## Supplemental file for "Astrocyte aquaporin mediates a tonic water efflux maintaining brain homeostasis"

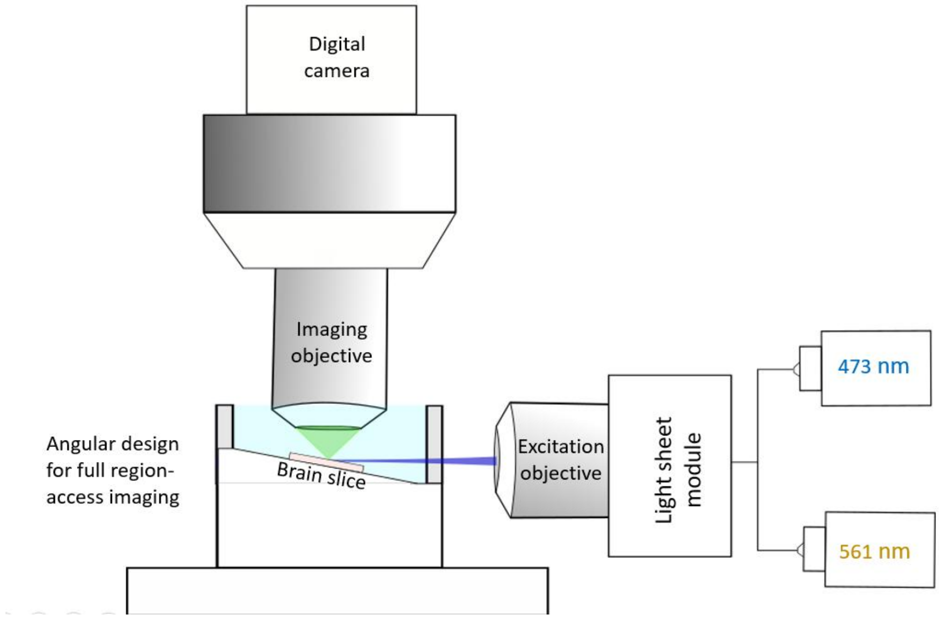


**Fig. S1.** Angular light sheet imaging in acute brain slices. Brain slice was placed at a slightly tilted (~8-12°) holding chamber so as to ensure the access of the light sheet to any regions of the slice, without interfering with the imaging objective. The light sheet illumination provides wide-field optical sectioning, while fluorescent images are collected by a digital camera via an independent imaging objective. See details in **Materials and methods**.

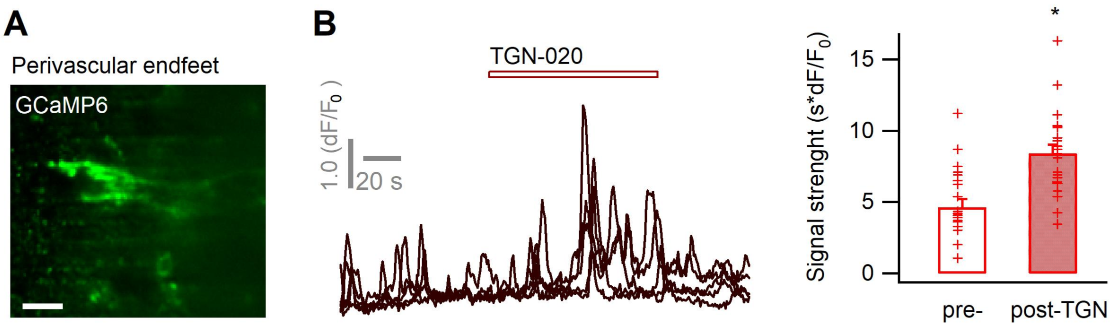


**Fig. S2.** (**A**) Imaging signals in the perivascular end feet of GCaMP6-expressing astrocytes. Scale bar, 20 µm. (**B**) The application of TGN-020 induced signal changes in the perivascular end feet (20 µM; n = 22 end feet ROIs from 3 mice). On average, the strength of basal $Ca^{2+}$signals in the end feet is higher than that observed across global astrocyte territories (4.65 ± 0.55 vs. 1.45 ± 0.79 Fig 2B, $p < 0.01$), as does the effect of TGN (8.4 ± 0.62 vs. 6.35 ± 0.97 Fig 2B, $p < 0.05$).

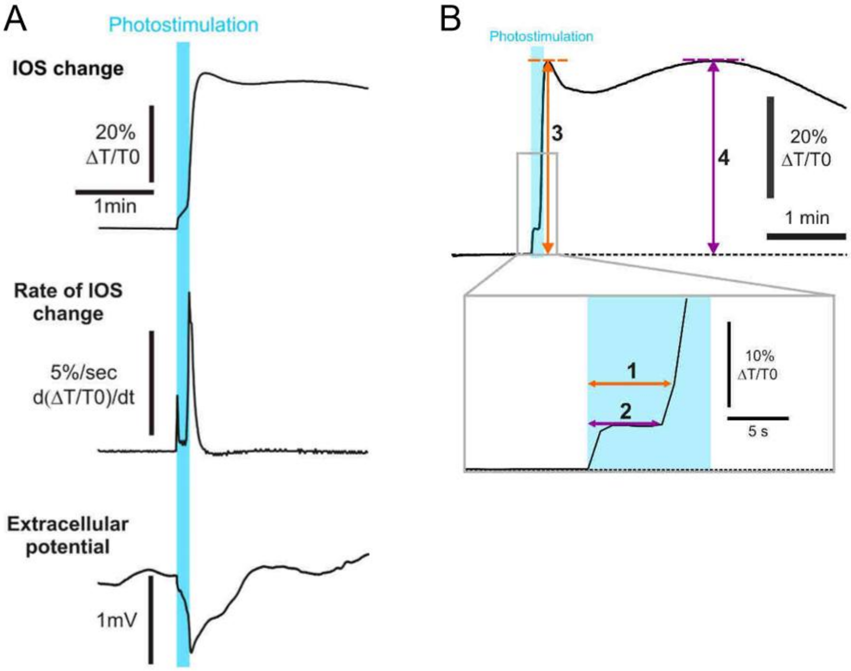


**Fig. S3.** Opto-CSD initiation and parameters analysis. (**A**) From the IOS change (*upper*), its temporal derivative was extracted to show the biphasic rate of the initiation of CSD (*middle*). Its starting, meanwhile, coincides with the extracellular potential drop (*bottom;* mean = -0.95 ± 0.21 mV, n = 4). (**B**) Parameter measurements for IOS changes: onset of the CSD (*1*) and the general swelling (*2*) relative to the beginning of the photostimulation, maximal IOS changes during CSD (*3*) and general swelling (*4*).

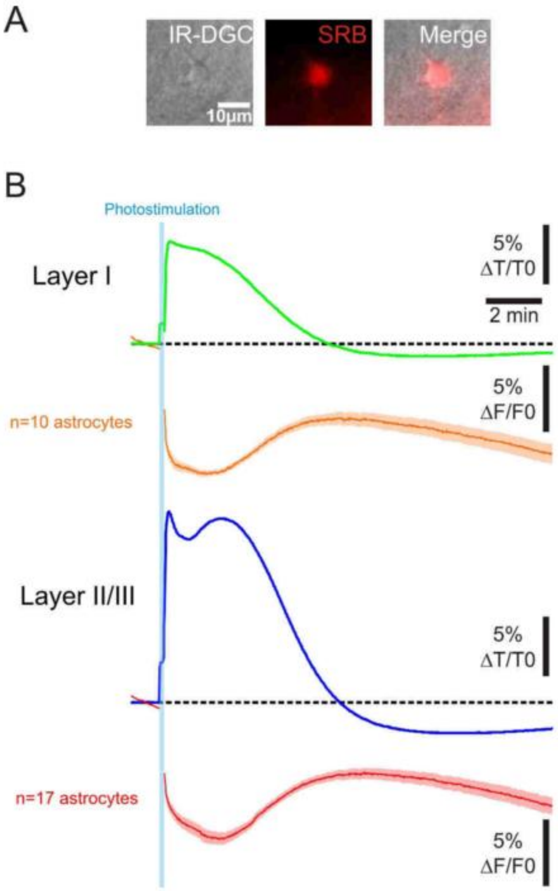


**Fig. S4.** Astrocytic swelling during an Opto-CSD across cortical layers. (**A**) Representative astrocyte SRB labeling observed under IR-DGC and epifluorescence. (**B**) IOS response (*green and blue traces*) and astrocytes swelling, reflected by SRB fluorescence decrease (*orange and red traces*), co-imaged in acute cortical slices. The dotted lines represent baseline and fluorescence imaging in fluorescent intensity. Astrocyte swelling spans over both the initial CSD response and the subsequent general cellular swelling of the parenchyma.

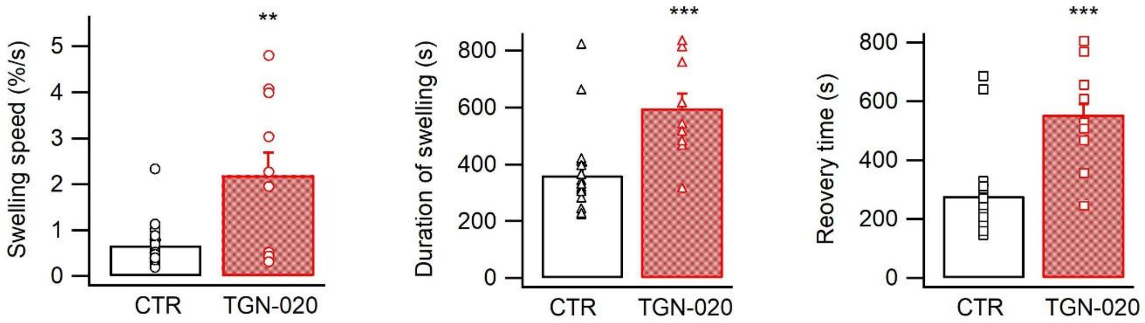


**Fig. S5.** The swelling speed, duration of swelling and recovery time during opto-CSD (n = 18 measurements from five mice for CTR, 10 measurements from four mice for TGN-020 for each parameter).

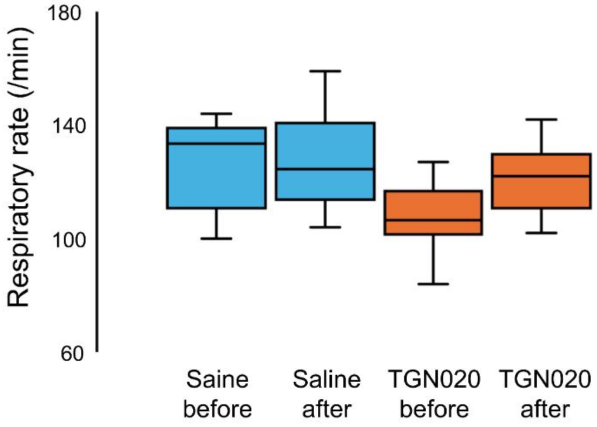


**Fig. S6.** Box-car plot of averaged respiratory rate before (0-10 min before injection) and after (50-60 min after injection) injection of saline or TGN-020.

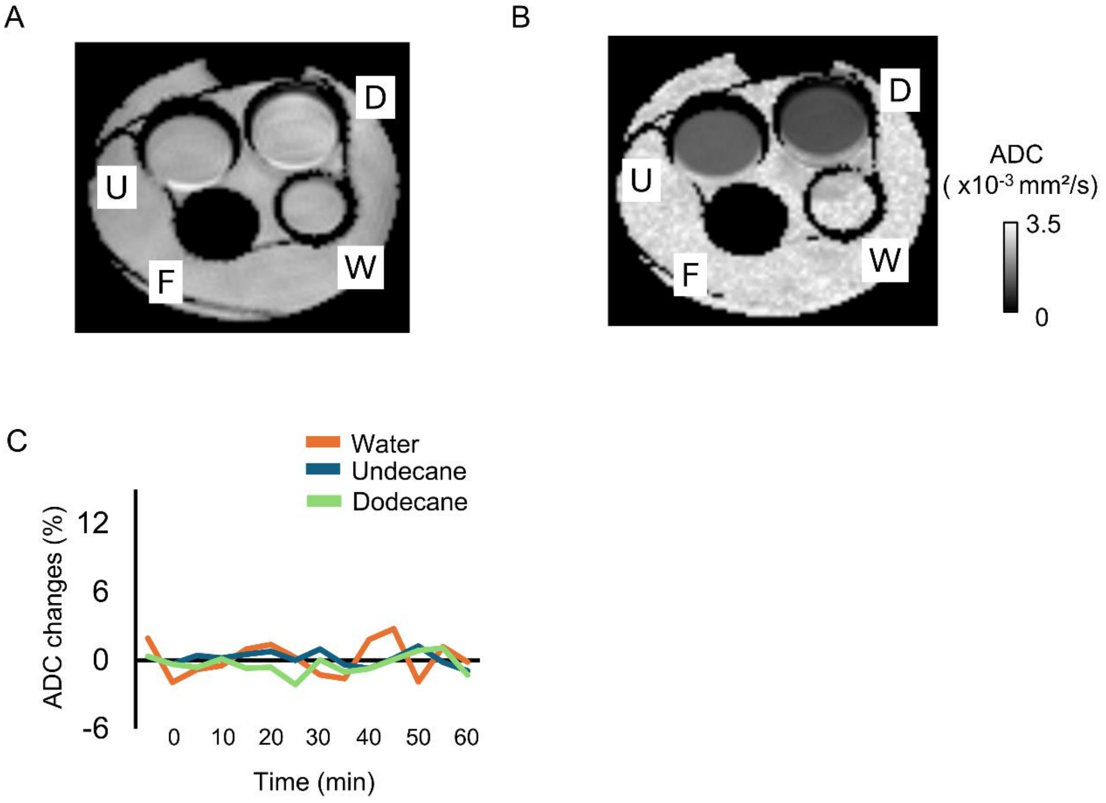


**Fig. S7.** Apparent diffusion coefficient (ADC) in water, undecane, dodecane, and fluorine (proton-free liquid). (A, B) Representative image obtained of (A) b = 0 s/mm² and (B) ADC map. D, n-dodecane; F, fluorine; U, undecane; W, water. (C) Time courses of water diffusion change in each solution was mapped by the calculation of ADC.

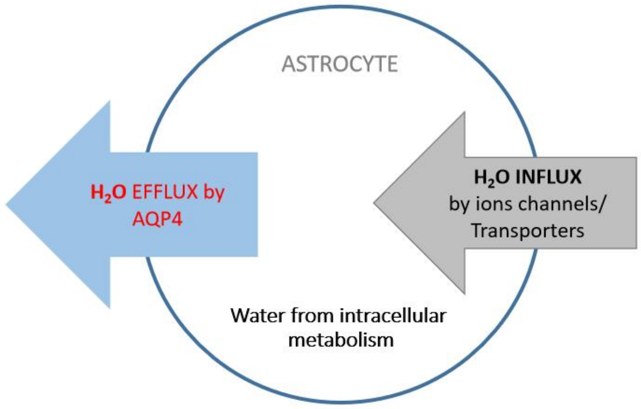


**Fig. S8.** AQP4-mediated tonic efflux helps to counterbalance excessive water accumulation in astrocytes, thereby maintaining water, volume and signaling homeostasis in the brain.
